# Pre-Supplementary Motor Area Theta Burst Stimulation Alters Corticomotor Facilitation and Action Reinitiation Without Impairing Response Inhibition

**DOI:** 10.64898/2026.08.10.743855

**Authors:** Eline Oen Lie, Aleksander Hagen Erga, Hayley J. MacDonald

**Author notes:** Corresponding author Jonas Lies vei 91 5009 Bergen Norway.

## Abstract

**Background:** The pre-supplementary motor area (preSMA) is increasingly being explored as a neuromodulation target for impulsive behaviour in several clinical populations. Treatment effects are generally interpreted as improvements in inhibitory control. However, healthy studies report improved/impaired/unchanged inhibitory control following identical preSMA stimulation protocols, and few studies examine accompanying neurophysiological changes. We therefore investigated whether preSMA stimulation influences downstream corticomotor excitability to modify a general stopping mechanism, other components of action control, or wider cue-dependent attentional processes relevant to impulsive behaviour.

**Methods:** In a preregistered, double-blind crossover study, 18 healthy adults received active and sham continuous theta burst stimulation (cTBS) over right preSMA. Motor-evoked potentials (MEPs), anticipatory response inhibition task measures, and alcohol dot-probe reaction times were collected before and after stimulation and analysed with linear mixed models.

**Results:** MEPs increased during sham (p = .028) but not after active cTBS (p = .741). Active cTBS did not affect complete or partial stopping on the response inhibition task. Instead, active cTBS slowed the continuing response after partial stopping (p < .001) whereas response execution sped up across the sham session (p < .001). No alcohol attentional bias or stimulation effect was detected.

**Conclusions:** PreSMA cTBS did not impair general inhibitory or attentional control. Instead, it attenuated session-related corticomotor facilitation and selectively slowed reinitiation of a partially inhibited action. These findings suggest that clinical effects to impulsive behaviour from preSMA neuromodulation are primarily rooted in changes to motor preparation and action updating rather than a unitary stopping mechanism.

## Introduction

Impulsivity is a multidimensional construct encompassing premature action, failures to suppress or revise prepared responses, and choices that favour immediate over delayed rewards [1, 2]. Disruption of these processes contributes to impulsive behaviours in neuropsychological conditions such as substance-use and behavioural addictions and impulse control disorders associated with dopaminergic treatment in Parkinson’s disease. Non-invasive brain stimulation (NIBS) has been proposed as a means of modifying the neural systems underlying impulsive behaviour in these clinical groups. However, addiction studies have predominantly targeted lateral prefrontal regions, particularly the dorsolateral prefrontal cortex, and have focused mainly on craving and clinical severity. Although the findings provide preliminary evidence of therapeutic benefit, variation in stimulation protocols, clinical populations, cognitive states and outcome measures has limited conclusions about the mechanisms involved [3, 4].

The pre-supplementary motor area (preSMA) has more recently emerged as a potential neuromodulation target because of its upstream position within fronto-basal-ganglia networks involved in selecting, stopping and revising actions. The preSMA is not a final motor output structure; rather, it influences whether prepared actions are initiated, withheld or replaced through interactions with the inferior frontal cortex, striatum, subthalamic nucleus and cortical motor regions. Paired-pulse transcranial magnetic stimulation (TMS) has demonstrated task-dependent influences of the preSMA on primary motor cortex (M1) during action reprogramming, while combined stimulation and neuroimaging studies indicate that preSMA modulation can alter activity and connectivity within preSMA–striatal–pallidal control circuits [5–8]. Motor-evoked potentials (MEPs) recorded from M1 may therefore provide a physiological measure of how an intervention applied to this upstream inhibitory control region propagates through the motor system.

Preliminary clinical findings support the preSMA as a candidate target but do not establish the cognitive or physiological mechanisms underlying treatment effects. Repeated preSMA continuous theta burst stimulation (cTBS) has been associated with reduced craving and symptom severity in gambling disorder and substance use disorders, and with improved inhibitory control performance in gambling disorder [9, 10]. Intermittent TBS over the preSMA has also improved response inhibition during sexual-cue exposure in medicated patients with Parkinson’s disease and hypersexuality, who showed altered preSMA–caudate connectivity [11]. These effects have generally been interpreted as improvements in inhibitory control. However, studies in healthy participants have reported both improved and impaired inhibition following preSMA cTBS [12, 13], and few studies have assessed whether behavioural changes are accompanied by altered corticomotor excitability. It therefore remains unclear whether preSMA stimulation modifies a general stopping mechanism, other components of action control, or wider cue-dependent processes relevant to impulsive behaviour.

The anticipatory response-inhibition task (ARIT) is well suited to separating these possibilities. Unlike tasks that provide only a single general measure of inhibition, the ARIT requires participants to prepare temporally predictable bimanual responses that must sometimes be cancelled completely or partially. Its outcome measures can distinguish anticipatory motor preparation, non-selective stopping, selective cancellation of one component of an action, and the updating or reinitiation of the continuing response following partial inhibition [14–17]. The ARIT may therefore identify more specific action-control processes through which preSMA stimulation could influence impulsive behaviour.

Attentional prioritisation of motivationally salient cues represents a related but distinct mechanism relevant to impulsivity in addiction and Parkinson’s disease-related impulse control disorders. For example, alcohol-related stimuli can preferentially capture attention, potentially increasing cue-induced motivation and craving. This attentional bias is commonly assessed using the alcohol dot-probe task (ADPT), in which faster responses to probes replacing alcohol-related rather than neutral images indicate preferential attention to alcohol cues [18, 19]. Whether preSMA stimulation affects this cue-biased attention, rather than motor preparation and inhibition alone, remains unclear. Although not principally an attentional-control region, the preSMA also contributes to resolving competition between automatically triggered and goal-directed responses and switching towards goal-appropriate behaviour [20]. The ADPT therefore provides a complementary test of whether preSMA stimulation alters the behavioural consequences of competition between salient cues and task goals.

The present sham-controlled study therefore examined the immediate effects of preSMA cTBS at three related mechanistic levels in healthy participants: 1) MEPs were used to assess changes in the downstream state of the corticomotor system; 2) the ARIT assessed action preparation, complete and partial stopping, and the reinitiation of partially inhibited responses; and 3) an ADPT assessed whether stimulation effects extended beyond action control to the attentional prioritisation of alcohol-related cues. Under a simple disinhibition account, we hypothesised that cTBS would reduce preSMA control over the motor system, increasing M1 excitability and impairing response inhibition, indexed by larger MEPs and longer stop-signal reaction times on the ARIT. We further hypothesised that active cTBS would increase alcohol attentional bias, expressed as a larger reaction time difference between probes following neutral versus alcoholic images on the ADPT. No pre-to-post changes were expected in any measures following sham stimulation.

## Methods

### Participants

Eighteen healthy participants were included in the study (3 male, 2 left-handed), aged 19-34 years (M = 24.5, SD = 3.93). Participants between 18-60 were recruited from the University of Bergen and through word-of-mouth. Participants fulfilled safety guidelines for TMS [21, 22], had no history of neuropsychological conditions, no current medications affecting dopamine and normal or corrected to normal vision. All participants gave their informed consent and the study was approved by Regionale Etiske Komiteer for medisinsk og helsefaglig forskning (REK; reference 774383). Monetary compensation was offered for participation (300kr).

### Design

The study was pre-registered (https://doi.org/10.17605/OSF.IO/8A69E), double blinded, and sham-controlled. Participants underwent two experimental sessions at least 6 days apart where they received active and sham cTBS (correctly identified post-study in 72% of participants) over the right preSMA in a counterbalanced order. Dominant M1 corticomotor excitability and behavioural measures from the ARIT and ADPT were recorded pre- and post cTBS. Data collection and analyses were as pre-registered, apart from switching to linear mixed models (LMMs) rather than Bayesian repeated measures ANOVAs as they better matched the structure of the final dataset. Importantly, the fixed effects tested in the LMMs corresponded to the preregistered experimental contrasts of interest. LMMs allowed us to account for participant-level differences in the size of condition effects, which is especially important given the variabiilty of responses to TBS [23].

### TMS procedures

A MagVenture R30 stimulator (MagVenture, Alpharetta, GA) with either an active (MagVenture MFC-B65) or sham (MagVenture MFC-P-B65) figure-of-eight coil was used. Two recording EMG electrodes (Ambu Neuroline 720 Neurology Surface Electrodes, recording system BLS pro) were placed in a belly-tendon montage on the dominant first dorsal interosseous (FDI) muscle. One grounding and one shielding electrode were placed on bony areas on the back of the hand/wrist. Single-pulse suprathreshold TMS was used for hotspotting to locate the dominant FDI muscle representation. The coil was held tangentially to the head with the handle at 45 degrees to the midline. The hotspot was marked with a washable pen on the scalp. Resting motor threshold (RMT) was recorded as the lowest stimulation intensity that obtained MEPs of at least 0.05 mV in 5 out of 10 consecutive trials. 15 MEPs were recorded at 120% RMT as a measure of corticomotor excitability at rest pre- and post TBS. Active motor threshold (AMT) was measured while participants held a constant weight (60 g padlock) on the end of their outstretched index finger to standardize level of activation (10% of maximum voluntary contraction). AMT was recorded as the lowest stimulation intensity that elicited MEPs of at least 0.2 mV in 5 out of 10 consecutive trials. cTBS was set to 80% AMT [12].

The right preSMA was marked 5 cm anterior to the vertex [24, 25] and 0.5 cm off the midline. For cTBS the coil was held at 0 degrees to the midline [26]. The cTBS protocol delivered three pulses at 50 Hz every 200 ms for 40 s (600 pulses total).

### Anticipatory Response Inhibition Task (ARIT)

The ARIT is computer-based and was performed in MATLAB (Version R2021a, MathWorks). Participants practiced each trial type in the first session to ensure thorough understanding of the task. Each trial started with the presentation of two vertical white rectangles that were intersected by a stationary target line near the top (Figure 1). When participants depressed and held down both the ‘Z’ and ‘-ߣ keys on the keyboard, two black bars started rising within the rectangles. The bars took one second to reach the top of the rectangles, while the target line was at 800ms. On Go trials (180 trials, 66%), the aim was to intercept the target line with the rising bars by lifting both fingers from the keys (Figure 1, A). Lift time (LT) was recorded relative to trial onset. On StopBoth trials (30 trials), the bars were programmed to stop rising before reaching the target, signaling participants must cancel their lift responses and keep both fingers on the keys (Figure 1, B). During Stop One trials, either the left (StopLeft; 30 trials) or right bar (StopRight; 30 trials) stopped before reaching the target, while the other bar continued rising (Figure 1, C and D). For a successful trial, participants had to keep down the finger corresponding to the stopped bar, while lifting their finger corresponding to the rising bar. Following established practice, each Stop trial type was individually staircased with an adaptive algorithm to convergence towards 50% success rate for each participant. For every correct stopping response, the bar stopped 25 ms later for the next Stop trial, and 25 ms earlier after each incorrect response.

**Figure 1.**
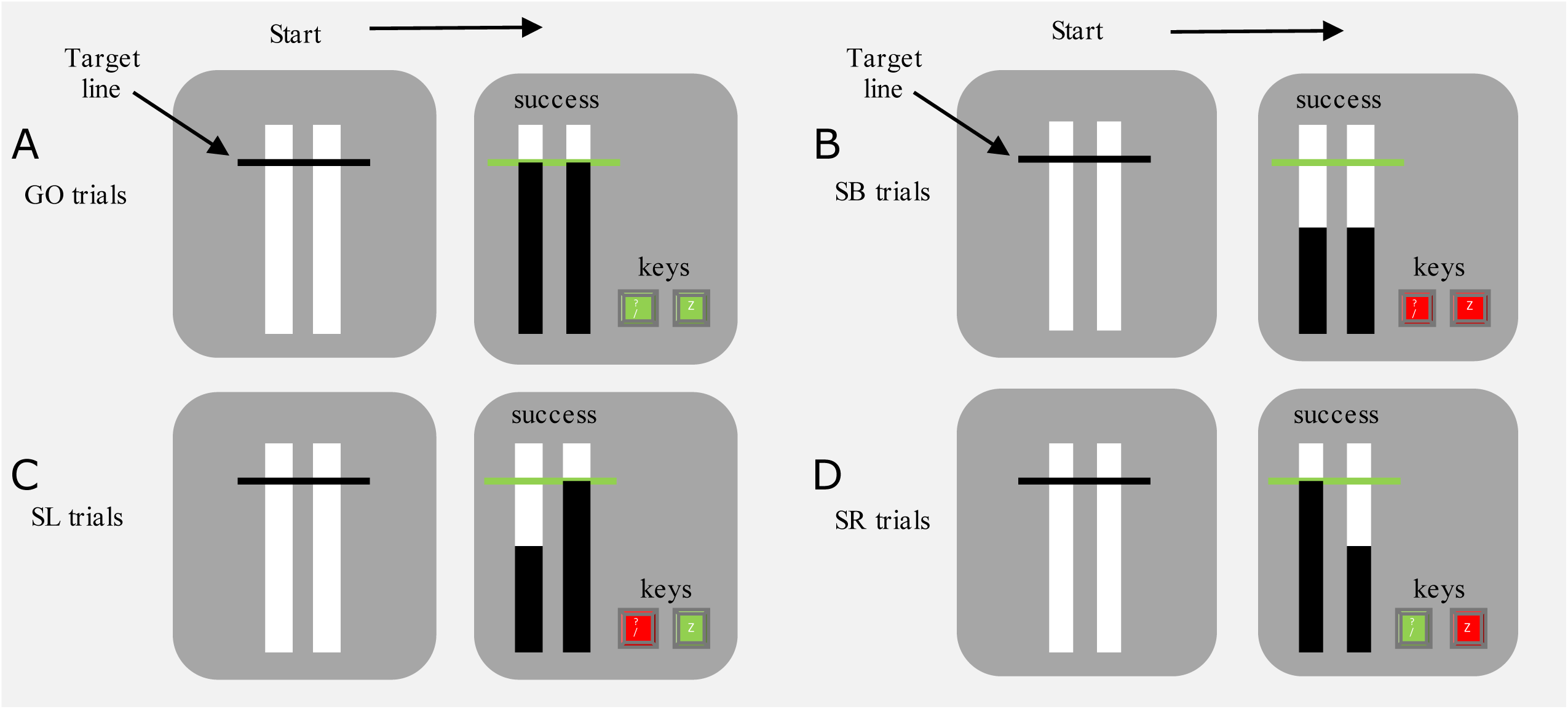
Trial types in the Anticipatory Response Inhibition Task. In the beginning, trial type was ambiguous. Green indicates the finger was lifted within 30 ms of the target line, red indicates the finger correctly remained on the key. A. Successful Go: both fingers are lifted at the target line. B. Successful Stop Both (SB): both fingers stay down when the bars stop rising before the target. C. Successful Stop Left (SL): the left finger stays pressing the key, the right is lifted at the target. D. Successful Stop Right (SR): the right finger stays pressing the key, the left is lifted at the target. Adapted from [46].

### Alcohol Dot Probe Task (ADPT)

The ADPT is computer-based and was performed using Inquisit 5. The script was adapted to Norwegian and images replaced with brands common in Norway to ensure recognition of beverages. The task started with a fixation cross in the middle of the screen presented for 500 ms. Subsequently two images were shown side by side for 1000 ms. The images presented were either one alcoholic and one non-alcoholic beverage, or a pair of non-beverage filler images. When the images disappeared, the probe (an X) appeared for 1000 ms in place of one of the alcoholic, non-alcoholic or non-beverage images on the left/right side of the screen (Figure 2). The task was to press the key as fast as possible indicating which side the probe appeared; ‘É for left, and ’Í for right. Images (matched on complexity) consisted of 10 alcoholic beverages, 10 non-alcoholic beverages, and 20 non-beverage filler images (10 pairs), presented 4 times each (80 trials total).

**Figure 2.**
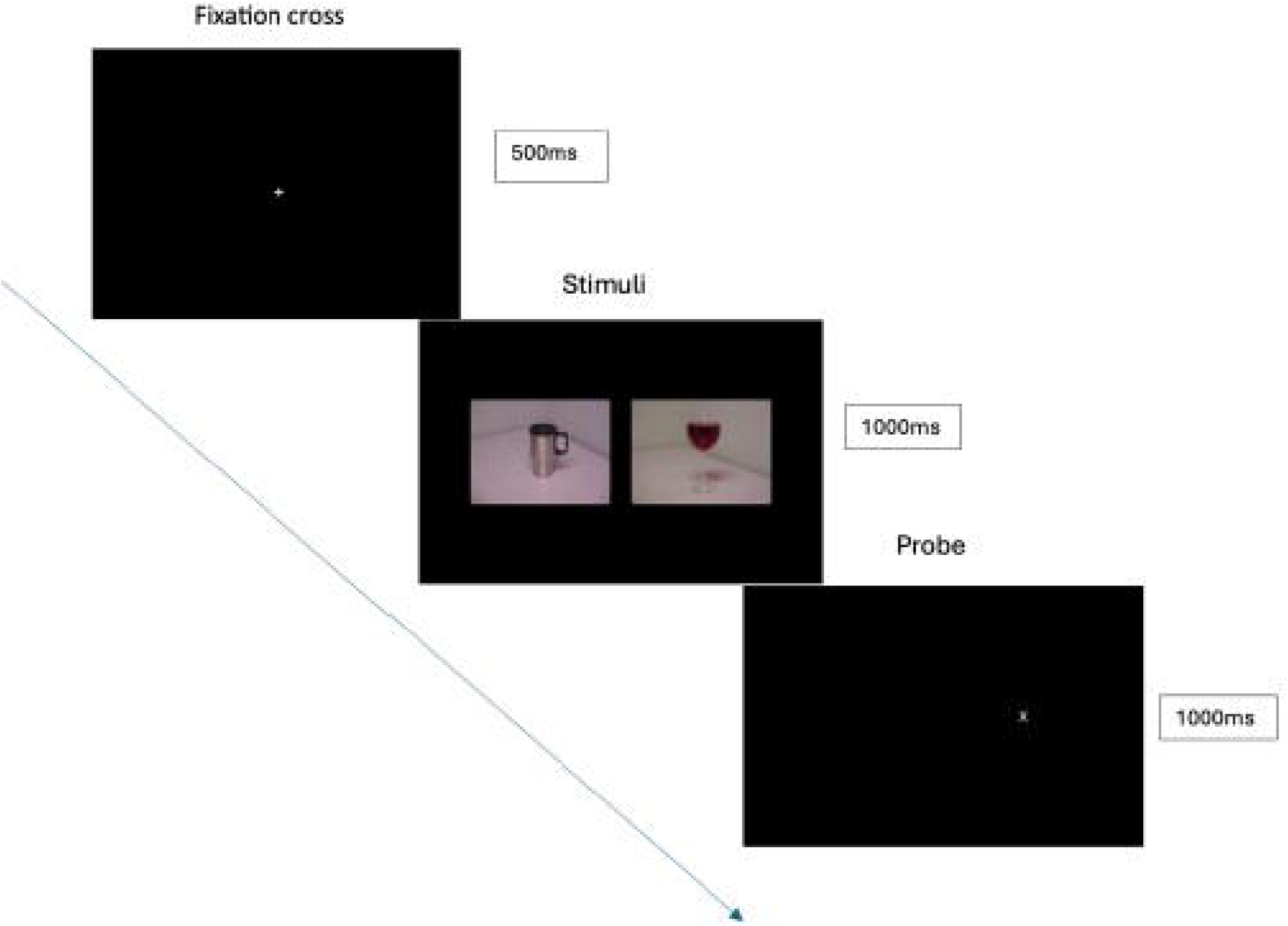
Trial progression in the Alcohol Dot Probe Task. A fixation cross appeared for 500ms, followed by an alcoholic/non-alcoholic beverage pair (shown) or filler images (not shown) for 1000ms. A probe (X) was subsequently presented for 1000 ms in place of one of the images and participants pressed the corresponding key (green) to indicate which side of the screen the probe appeared.

### Data analysis

Primary dependent neurophysiological measures were pre- and post TBS M1 excitability (peak-to-peak MEP amplitude), processed using BSL Analysis (version 4.1). Root mean square EMG (rmsEMG) was recorded 5-55 ms prior to the TMS pulse as a secondary measure, and cut off for MEP inclusion was 0.02 mV.

Primary dependent measures for the ARIT were pre- and post TBS stop signal reaction time (SSRT) on successful StopBoth, StopRight and StopLeft trials. SSRT was calculated using the integration method [27]. Secondary measures were LT (after trimming outliers ± 3 SD) [14, 28] on Go, StopLeft and StopRight trials. Trials without a LT were removed from the analysis. Primary dependent measures for the ADPT were pre- and post TBS reaction time (RT) on correct trials when the probe replaced alcoholic and non-alcoholic images. RT was measured from when the probe appeared until a key was pressed. The secondary measure of RT on non-beverage filler trials served as a control condition.

### Statistical analyses

R (Version 4.4.1) was used to conduct LMMs (*lmer*) on all primary and secondary dependent measures. All LMMs included fixed effects of Time (Pre, Post) and Stimulation Type (Active, Sham) with their interaction, Participant as a random effect, and a random slope for Time. The LT LMM included an additional fixed effect of Trial Type (Go, StopOne) and its interaction with Time and Stimulation Type. An error in the code meant only the first LT of the two fingers on Go trials was reliably recorded, so Side (Left, Right) was not included in the model. RTs included an additional fixed effect of Trial Type (Alcoholic, Non-alcoholic) with its interactions, and a random effect of Image. Normality was assessed by Shapiro-Wilk and Q-Q plots of residuals in JASP (Version 0.19.3). MEP and RT data violated assumptions of normality, and were log transformed in the LMMs. Post-hoc comparisons were performed where necessary using *emmeans* and pairwise Tukey corrections for multiple comparisons, with statistical significance set at α=0.05. Results are reported as mean ± standard error.

## Results

Two participants were excluded from StopOne trial analyses due to an inability to adequately perform these trials in both sessions. SSRT and LT models including StopOne trial data therefore include N = 16. For all other measures N = 18.

### TMS measures

There was an increase in M1 excitability across the sham session that was absent after active stimulation. As predicted, MEP amplitude showed a significant interaction effect (E = −0.421, p = .038) but contrary to our hypothesis, it was sham stimulation that produced a significant increase in MEP amplitude from pre (0.582 ± 0.112 mV) to post (0.861 ± 0.200 mV, p = .028), while active stimulation did not (pre: 0.623 ± 0.112 mV; post: 0.750 ± 0.218 mV; p = 0.741; Figure 3). Furthermore, baseline (pre) MEP amplitude in the sham session was significantly lower than in the active cTBS session (p < 0.001). There was no main effect of Stimulation Type (E = −0.057, p = .874) or Time (E = 0.039, p = .813). Paired sample t-tests showed no significant difference in RMT (p = 0.448) or AMT (p = 0.453) between sessions, indicating a change in stimulation intensity could not account for baseline differences in MEP amplitude or effects of cTBS.

**Figure 3.**
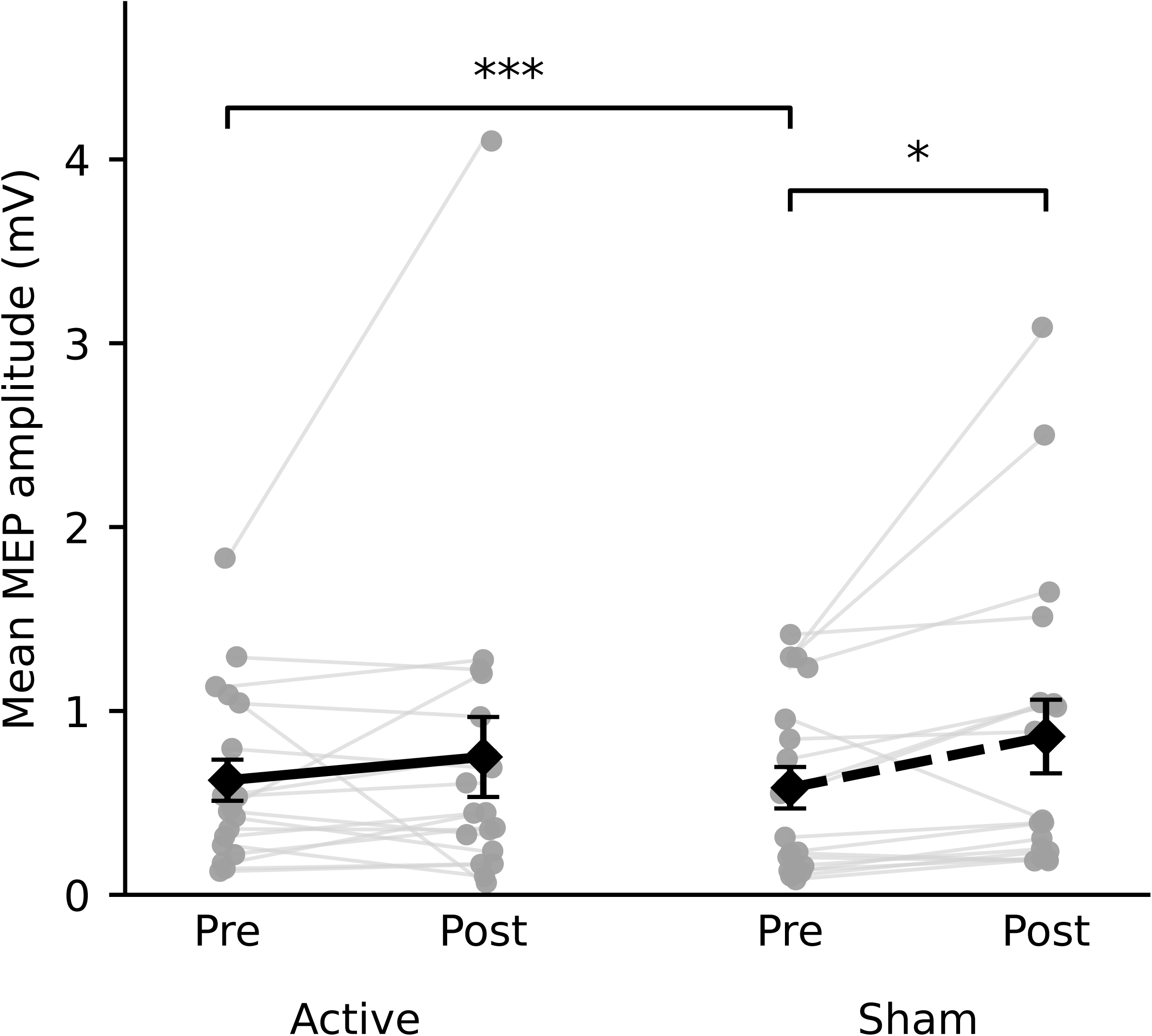
Corticomotor excitability results. Raw motor evoked potential (MEP) amplitude increased across the sham session (dashed line, *p<0.05) but no difference was seen from pre- to post active stimulation (full line). Baseline (pre) MEP amplitude was lower in the sham compared to active sessions (***p<0.001). mV: millivolts.

Differences in corticomotor excitability were not driven by background muscle activity. For rmsEMG, there was a main effect of Time (E = 0.0008, p = .006). Collapsed across Stimulation Type, pretrigger muscle activity at rest decreased from pre (0.007 ± 0.003 mV) to post (0.006 ± 0.003 mV) stimulation. There was no main effect of Stimulation Type (E = 0.0003, p = .370), and importantly no interaction effect (E = −0.0005, p = .259) that could explain the MEP amplitude results.

### ARIT

Contrary to our hypothesis, active cTBS had no effect on general inhibitory control of motor behaviour. StopBoth SSRT showed no main effect of Stimulation Type (E = 0.014, p = .058) or Time (E = −0.003, p = .675) and no interaction effect (E = −0.006, p = .582). StopOne trial SSRTS also showed no main effect of Stimulation Type (StopLeft: E = −0.003, p = .812; StopRight: E = −0.009, p = .402) or Time (StopLeft: E = −0.001, p = .931; StopRight: E = −0.012, p = .275), and no interaction effects (StopLeft: E = 0.005, p = .800; StopRight: E = 0.013, p = .382).

Active cTBS slowed selective response reinitiation following complete cancellation, while basic response execution sped up across the sham session. For LTs on Go and StopOne trials, there was a Trial Type x Stimulation Type x Time interaction (E = 0.013, p = 0.002; Figure 4) which showed that Go trial LTs decreased from pre (815 ± 3 ms) to post (808 ± 2 ms, p < 0.001) in the sham condition, and active stimulation produced an increase in StopOne LTs from pre (878 ± 3 ms) to post (896 ± 2 ms, p < 0.001). There was the expected main effect of Trial Type (E = 0.072, p < 0.001) with a robust delay on StopOne (887 ± 2 ms) compared to Go trials (812 ± 2 ms) that confirmed the task was performed as expected.

**Figure 4.**
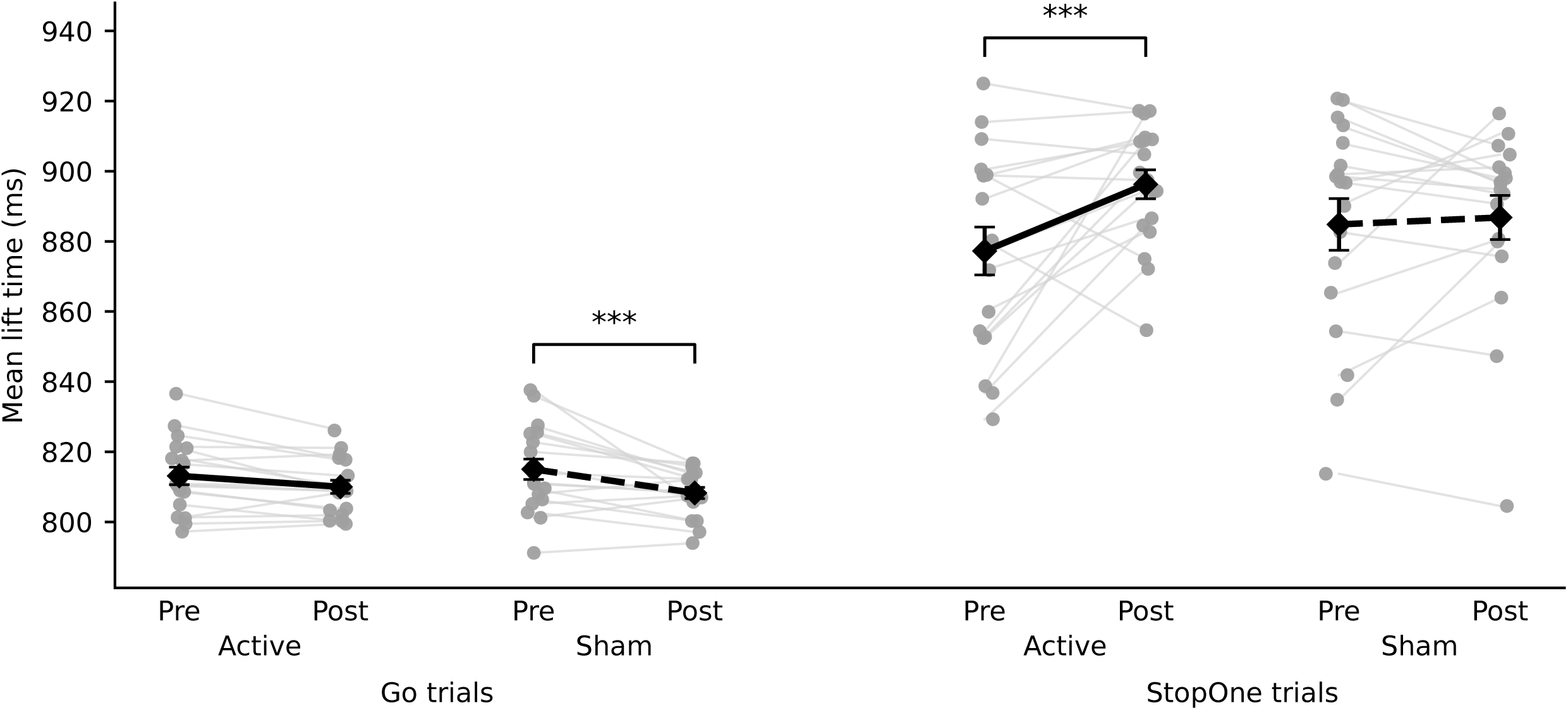
Lift times on Go and StopOne trials of the Anticipatory Response Inhibition Task. Go trial lift times (reported in milliseconds, ms) decreased across the sham session (dashed line), while StopOne lift times increased from pre to post active stimulation (full line). ***p<0.001.

### ADPT

Contrary to our hypothesis, active cTBS did not affect the ability to suppress or disengage from alcohol-related cues. Surprisingly, there was no attentional bias regarding alcoholic images in our participants. RTs on trials with alcoholic versus non-alcoholic beverage images showed no main effect of Trial Type (E = 0.003, p = 0.755), Stimulation Type (E = 0.010, p = 0.743) or Time (E = 0.003, p = 0.326) and no interactions (all p > 0.349). An effect of Trial Type would reflect attentional bias towards/against alcoholic images. Filler image RTs also showed no significant effects (all p > 0.291), indicating no changes in general RT.

## Discussion

The current study investigated the effect of sham-controlled cTBS over the right preSMA at three mechanistic levels: downstream corticomotor excitability, action control, and alcohol-cue-biased attention. Contrary to our hypothesis, active preSMA cTBS did not increase resting M1 excitability and impair inhibitory or attentional control. Instead, the general pattern of results suggests that there was a nonspecific increase in motor readiness or adaptive motor facilitation across the session, reflected in larger MEPs and faster Go lift times across sham stimulation. Active preSMA cTBS attenuated this facilitation and selectively disrupted the updating/reinitiation of a partially inhibited motor response, reflected by slower StopOne LTs. The absence of corresponding changes in filler-trial RTs on the ADPT argues against a general slowing of perceptual or motor responses. However, the lack of an alcohol-related attentional bias under either stimulation condition limits conclusions regarding the effects of preSMA cTBS on cue-biased attention. While background muscle activity and changes in stimulation intensity cannot account for corticomotor excitability results, it is important to bear in mind baseline MEP differences between active and sham sessions which might implicate a role of state-dependence on TMS aftereffects and replication of results with comparable baselines would be prudent. Nevertheless, the current convergent neurophysiological and behavioural findings suggest that preSMA cTBS did not produce a broad impairment of inhibitory or attentional control but instead altered the facilitation and updating of prepared actions. These findings shift the mechanistic interpretation of preSMA NIBS away from a simple increase or decrease in inhibitory control and towards modulation of how interrupted or partially inhibited actions are subsequently reconfigured and implemented.

The direction of the neurophysiological effect was contrary to the original hypothesis. In line with traditional accounts, we predicted that reducing preSMA activity would weaken inhibitory control over the motor system and consequently increase M1 excitability. However, this prediction assumes that the preSMA exerts a predominantly inhibitory influence on M1. Evidence exists that its influence is task-and time-dependent. During action reprogramming, the preSMA facilitates the M1 representation of the newly appropriate response, whereas the right inferior frontal cortex exerts a later inhibitory influence on the response that must be cancelled; disruption of the preSMA also alters this right inferior frontal–M1 interaction [5, 29]. The preSMA may therefore coordinate response selection, threshold adjustment, and recruitment of the wider inhibitory network rather than directly suppressing corticomotor output [30]. Attenuation of the session-related MEP increase following active cTBS could consequently reflect reduced facilitatory or preparatory input to M1 rather than increased inhibition within M1. This interpretation is consistent with the selective slowing of StopOne lift times (with no corresponding effect on SSRTs), because these trials required participants to update and reinitiate a unimanual motor response after bimanual non-selective inhibition [14–16]. PreSMA disruption has similarly been associated with impaired updating of motor plans, and the region’s influence on M1 is particularly facilitatory when an alternative response must be implemented [29, 31]. The convergence of altered MEP amplitude and response reinitiation time, in the absence of an SSRT effect, may therefore suggest that the most evident consequence of cTBS occurred at the level of cortical motor preparation and implementation rather than through a general disruption of the cortico-basal-ganglia pathways supporting rapid stopping. One possible route is altered cortico-cortical influence from the preSMA through SMA proper or premotor regions to M1 [32]. This should not, however, be interpreted as evidence for an exclusively cortical mechanism: indirect striato-pallidal and hyperdirect cortico-subthalamic pathways interact during response inhibition and may also contribute to action selection and reinitiation [6, 33]. Nevertheless, because resting MEPs provide an indirect measure of preSMA network effects and cTBS cannot be assumed to produce uniform local inhibition outside M1, this mechanistic interpretation needs further verification.

There are, however, some alternative or contributing explanations for the current results apart from the central mechanism presented so far. Sham MEP amplitude was lower than active MEP amplitude at baseline despite comparable background muscle activity, indicating differences in resting cortical state. Baseline variability can predict part of the response to cTBS [34], while TBS after-effects vary across individuals according to the interneuronal circuits recruited and the ongoing cortical state [23, 35]. Consistent with a nonspecific session effect, sham-controlled cTBS research has reported increases in MEP amplitude over time irrespective of stimulation condition [36]. However, such facilitation could potentially reflect arousal, or stress rather than motor readiness: experimentally induced worry, psychosocial stress, and performance pressure can transiently increase MEP amplitude [37, 38]. Although the contribution of these effects remains speculative, they could be easily tested using session-specific anxiety and physiological arousal measures in future research.

At the third mechanistic level, active preSMA cTBS did not alter alcohol-cue-biased attention on the ADPT. This task indexes competition between cue-driven and goal-directed attention, but its conventional RT contrast cannot distinguish facilitated orienting from delayed disengagement or determine whether any bias reflects motivational salience, reduced top-down control, or both. More importantly, no alcohol-related attentional bias was detectable under either stimulation condition, leaving little behavioural expression for cTBS to modulate. Such biases are more consistently observed in heavier drinkers than in healthy social drinkers, although they can occur in non-problem-drinking university samples [18, 39–41]. The null ADPT result therefore provides limited evidence about whether preSMA NIBS can alter clinically relevant cue processing. Nevertheless, the absence of effects on both beverage and filler trials, alongside changes in MEP amplitude and ARIT lift times, suggests that the present effects were expressed more strongly in motor system state and action updating than in cue-biased attention. Although this pattern should not be taken as a definitive dissociation between action and attentional control because the tasks differ in sensitivity and psychometric properties.

Clinically, the effects of altered preSMA-centred circuitry may lie more in motor preparation and action updating than in a general reduction of inhibitory control or cue processing. In addiction and Parkinson’s disease-related impulse control disorders, behavioural impairments may therefore involve not only failure to suppress a prepotent action, but also difficulty reconfiguring and implementing the appropriate response after an action has been interrupted. This may help explain why clinical studies report changes in craving, symptom severity, or selected dimensions of impulsivity without consistent effects across all outcomes. The effects of preSMA NIBS are likely to depend on the component of action control measured, baseline neural state, and motivational context. Future studies should therefore combine process-specific measures of stopping, switching, and response reinitiation with physiological or network-level measures of target engagement.

Two limitations should be considered. First, the intended right preSMA target was only 0.5 cm lateral to the midline and was not individualised using structural MRI. The effects are therefore best interpreted as right-biased stimulation of medial preSMA, with possible bilateral engagement, as similarly acknowledged by Obeso et al. [7]. Second, nicotine use was not controlled and may have contributed to between-participant or between-session variability. Acute and chronic nicotine exposure can alter M1 inhibitory and facilitatory circuits, NIBS-induced plasticity, and attentional performance [42–45]. This may be relevant in Norwegian student samples where oral nicotine use is common. Future studies should record nicotine status or standardise recent exposure.

## Acknowledgements

The authors thank Andrea Bøe, Rebekka Hilton Fyllingen Granerud and Silje Moen for assisting in participant recruitment and data analysis.

## Declaration of generative AI and AI-assisted technologies in the manuscript preparation process

During the preparation of this work the authors used Open AI to help with brevity and clarity of writing. After using this tool/service, the authors reviewed and edited the content as needed and take full responsibility for the content of the published article.

## Data availability statement

The anonymised datasets analysed in the current study are available on GitHub (XXX).

